# A cellular and spatial map of salivary glands at single cell resolution reveals the functional basis of tertiary lymphoid structure formation in Sjogren’s syndrome

**DOI:** 10.1101/2022.11.03.514908

**Authors:** Saba Nayar, Jason D. Turner, Saba Asam, Eanna Fennell, Matthew Pugh, Serena Colfrancesco, Onorina Berardicurti, Charlotte G. Smith, Joe Flint, Ana Teodosio, Valentina Iannizzotto, David H. Gardner, Joel van Roon, Ilya Korsunsky, Roche Fibroblast Network Consortium, Simon J. Bowman, Wan-Fai Ng, Adam P Croft, Andrew Filer, Benjamin A. Fisher, Christopher D. Buckley, Francesca Barone

## Abstract

The key role of tertiary lymphoid structures in autoimmune and non-autoimmune conditions has been recently appreciated. While many of the molecular mechanisms involved in tertiary lymphoid structure (TLS) formation have been identified, their cellular sources and their temporal and spatial relationship to each other during the development of TLS remain unknown. Here we have constructed a cellular and functional map of key components involved in the formation of TLS in the minor salivary glands (SG) in humans. We have confirmed the presence of an immunofibroblast cell state and identified an undescribed immunopericyte cell state with potential immunological functions within TLS. The identification of TLS cellular and functional properties and their relevant modulators provided by this analysis provides key therapeutic cues for TLS associated conditions in autoimmunity and cancer.

## Introduction

Tertiary lymphoid structures (TLS) provide a functional microenvironment for activated T and B cells to reside, proliferate and differentiate within non lymphoid tissues^1, 2, 3, 4, 5, 6, 7, 8, 9^. Whilst unlikely to play a role in disease initiation, TLS support disease persistence and have been shown to associate with clinical features, response to therapy and progression in a range of autoimmune diseases and in cancer^10^. Associations with pathophysiology have been most clearly demonstrated in Sjögren’s syndrome (SjS), a disease where formation of TLS and maturation of ectopic germinal centers in lymphoid structures correlate with B cell hyperactivity, autoimmunity, pathogenic humoral response and development of B cell lymphoma^5, 11, 12, 13, 14, 15, 16, 17^.

We and others have previously demonstrated that FAP^+^PDPN^+^ immunofibroblasts, a population of tissue resident fibroblasts that share features and functions with fibroblastic reticular cells inhabiting secondary lymphoid organs (SLOs), define the anatomical framework for TLS establishment and maintenance in peripheral tissue^5, 13, 18, 19^. Analysis of murine models of TLS and human samples from patients with SjS has unveiled multiple signals and molecular mechanisms involved in the activation of immunofibroblasts, leading us to propose the presence of a PDGF-Rα^+^/PDGF-Rβ^+^ tissue resident progenitor cell that differentiates in response to local cues, to acquire novel immunological properties^5^. Inflammatory and lymphoid cytokines such as IL-13, IL-22 and both lymphotoxin alpha and beta (LTα3 and LTα1β2) have been shown to play a role in the activation and differentiation of immunofibroblasts^5^.

Use of computational biology approaches applied to single cell transcriptomic data has provided novel cellular atlases of healthy and disease tissues and molecular signatures to identify “gene cassettes”, enabling the categorization of fibroblast clusters that share similar features across different diseases and in different organs^20, 21^. Using cellular interaction inference tools within scRNAseq data, investigators have also been able to propose molecular interactions between specific cell populations. An example of these findings is the identification of the role of NOTCH3 in establishing a transcriptional gradient for fibroblast differentiation in the synovial tissue in Rheumatoid Arthritis^22^ and the discovery of the complement component C3 as a mediator in macrophage/cancer associated fibroblast (CAFs) crosstalk^23^. The complexity of the stromal cell compartment of murine lymph nodes (LN) has been also deconvoluted, unveiling an unexpected level of heterogeneity and cellular specialization within lymph nodes^24^. Different populations of fibroblasts create specialized micro domains within the LN, which influence antigen presentation, cell migration, retention, activation and survival of T and B lymphocytes^11, 25, 26, 27, 28, 29, 30^, all of which ultimately shape the immune response.

So far, human tissues fostering TLS structures have not been explored at single cell resolution, leaving an important gap in our understanding of the cellular and molecular properties involved in TLS establishment, maintenance, and evolution. Importantly, as for other complex dynamic microenvironments, TLS single cell analysis needs to be integrated within the context of TLS histological organization to provide a developmental reference for disease pathophysiology. Here we have determined the cellular and spatial basis of TLS formation in SjS to unveil TLS specific properties, identifying cellular players, delineating key pathways involved in cellular cross talk and defining the developmental trajectory of TLS formation and function in peripheral tissue. This study, whilst providing the first comprehensive map of TLS cellular and molecular properties, provides tools to promote understanding of these structures in disease and to select specific modulators for TLS manipulation in human disease.

## Results

### Single cell analysis unveils the complex cellular landscape of TLS in human salivary gland

To gain single cell resolution of the inflamed glandular microenvironment in salivary glands (SG) harboring TLS and to better understand cellular properties associated with ectopic lymphoneogenesis, we performed 10x Single-cell RNA sequencing on labial minor SG biopsies obtained from seven patients with SjS (Supplemental Data: Table S1; 17,011 cells passed QC) (fig 1a). Within the global cellular landscape of the human SjS SG, we identified 11 main cell clusters consisting of 5 hematopoietic and six stromal cell clusters. In the CD45^-^ stromal cell compartment we identified epithelial/myoepithelial, endothelial, fibroblast, mural, and a small neuronal cell population. Subclustering of the CD45^+^ hematopoietic compartment unveiled presence of plasma cells, B cells, CD4^+^ and CD8^+^ CD3^+^T lymphocytes, and myeloid cells (fig 1b). Multiplex immunofluorescence with differentiating markers for each of the identified cell types (fig 1c), combined with analysis of gene cassettes associated with each cell type (suppl. fig 1), confirmed our annotation of the hematopoietic and stromal populations. Cluster distribution was validated across all analyzed samples (suppl. fig 1).

**Figure 1:**
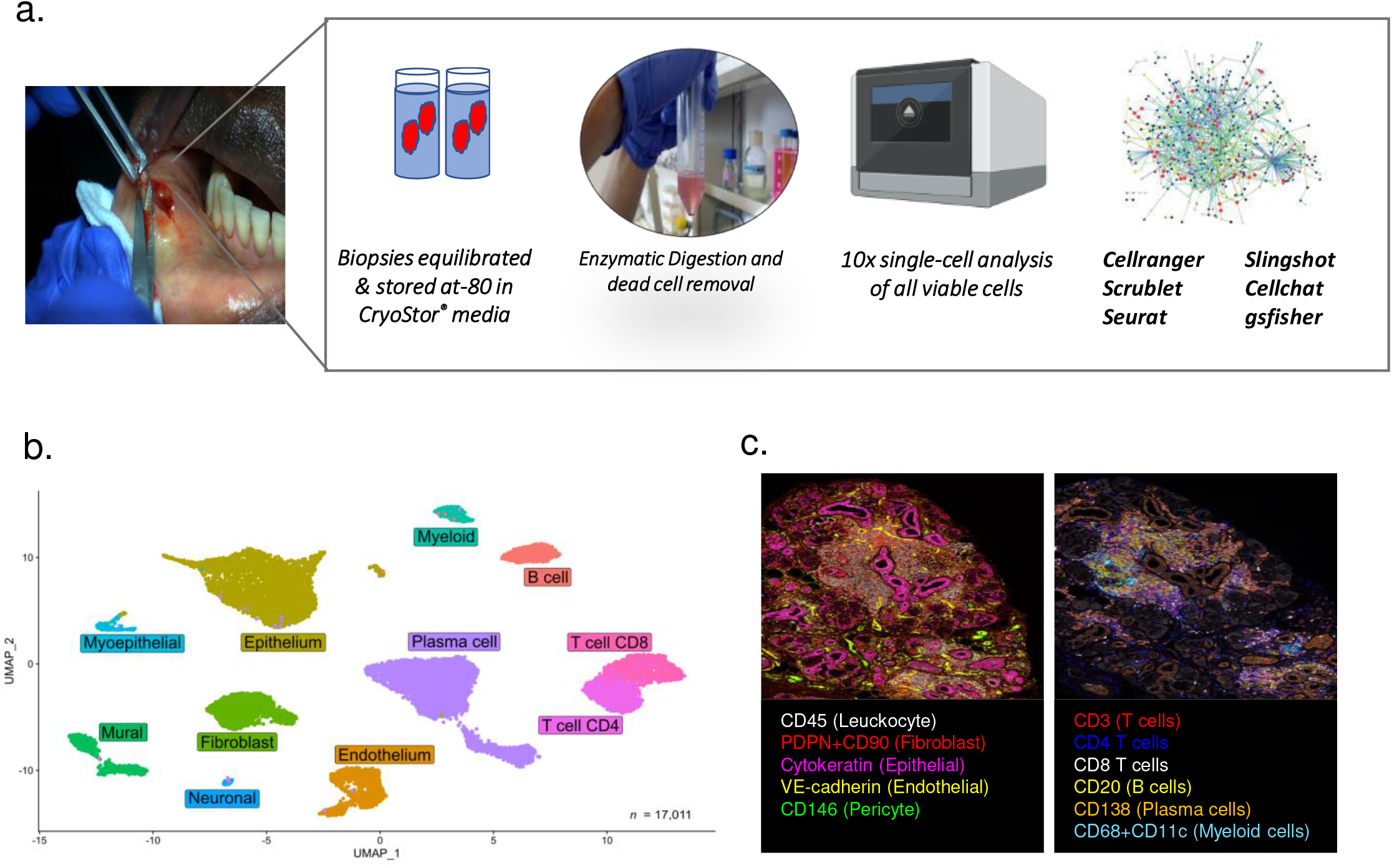
Cellular landscape of the minor salivary glands of SjS patients: **a**, Minor salivary gland biopsy 10x single cell workflow pipeline. **b**, Uniform manifold approximation and projection (UMAP) embedding of 17,011 single cells and gross-cell identities clustering of single-cell gene expression data from n=7 Sjogren’s syndrome salivary gland samples illustrating both immune and stromal cell clusters. **c**, Multiplex immunofluorescence image illustrating major lineages identified in minor salivary gland tissue probed with a 6-plex panels for CD146 (green), VE-cadherin or CD20 (yellow), Pan-Cytokeratin/CK (magenta), CD138 (orange), and CD68+CD11c (cyan), CD45 or CD8 (white), Podoplanin/PDPN+CD90 or CD3 (red) and CD4 (blue).

### Stromal cell clustering reveals significant fibroblast diversity and a novel immunopericyte population in Sjögren’s syndrome

We and others have previously demonstrated that fibroblasts are key in supporting TLS development and maintenance in the tissue^5, 10, 13, 23, 31, 32, 33, 34, 35^. However, heterogeneity is present in this compartment, that must be resolved, to develop the ability to manipulate TLS establishment. To this end, we performed sub-clustering on CD45^-^ cell clusters (fig 2a). Extensive differential expression analysis was implemented to validate the cellular identity of the annotated clusters (suppl. fig 2) and key differentiating genes, identified using the implementation of the MAST algorithm within Seurat, as summarized in fig 2b.

**Figure 2:**
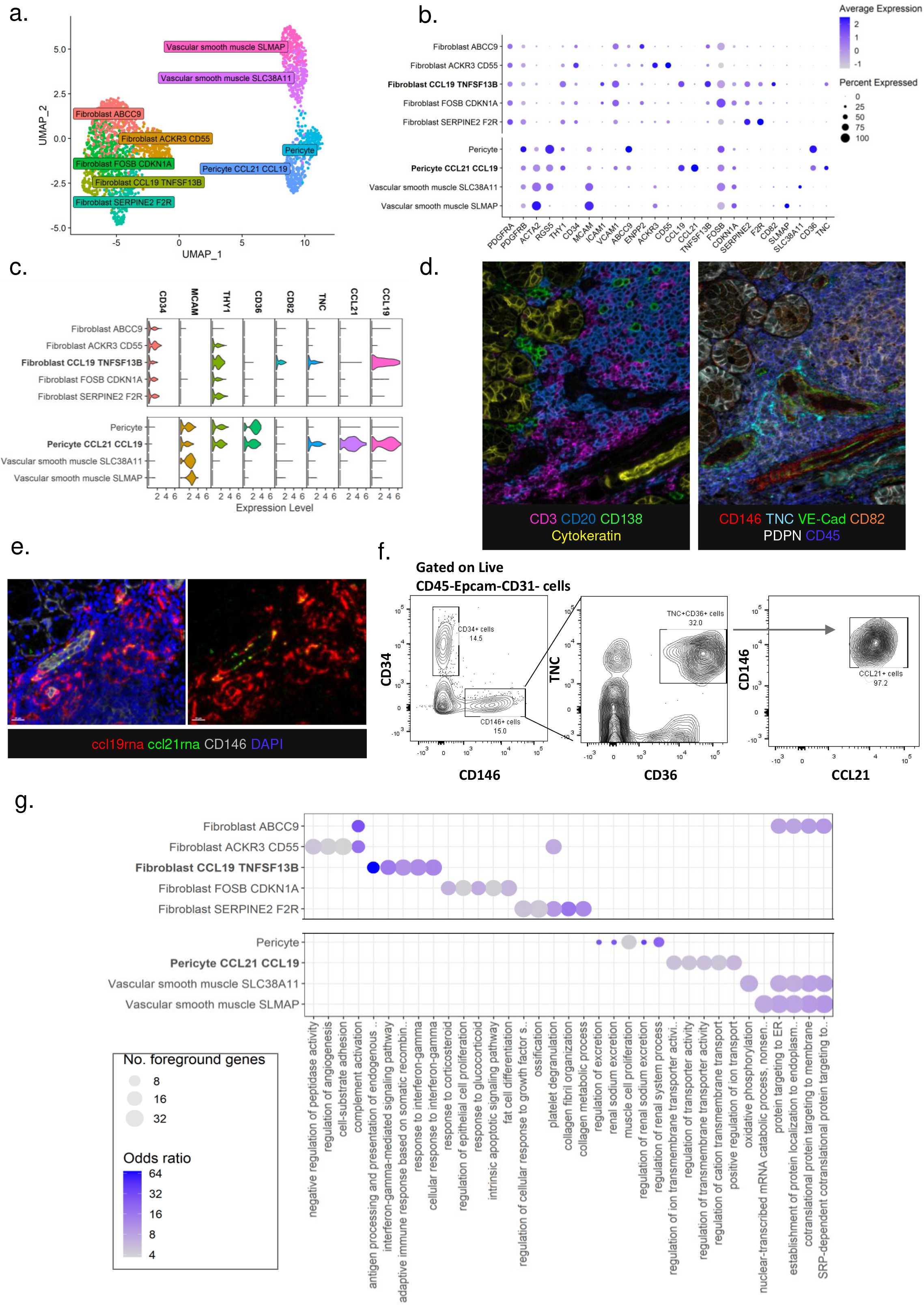
Single cell resolution of fibroblast and mural cell subsets in human SjS salivary glands: **a**, Uniform manifold approximation and projection (UMAP) dimensional reduction and sub-clustering of fibroblast and mural cells in Sjogren’s syndrome minor salivary glands. **b**, Expression of current identifying and newly discovered identifying genes across the fibroblast and mural subsets. **c**, Expression of gene cassettes used to spatially identify immunofibroblast and immunopericytes in salivary glands **d**, Multiplex immunofluorescence image illustrating immune cells and pericytes identified in minor salivary gland tissue probed with a 6-plex panels for CD138 (green), CD20 or TNC (yellow), Pan-Cytokeratin/CK or CD82 (orange), VE-cadherin (cyan), Podoplanin/PDPN (white), CD3 or CD146 (red) and DAPI or CD45 (blue). **e**, Multiplex immunofluorescence image of CD146 (grey) pericytes with ccl21rna (green), ccl19rna (red) and DAPI (blue) in immune aggregate within human salivary glands of patients with Sjögren’s syndrome. **f**, Representative flow cytometric identification of CD146+TNC+CD36+CCL21+ immunopericyte cell population in Sjogren’s syndrome minor salivary glands. **g**, Gene set enrichment analysis (using GO and Kegg gene sets) of the fibroblast and mural populations, the top 5 enriched gene sets for each cell subset are shown.

Immunofibroblasts have been previously described both in humans and mice as CD45^-^ cells expressing ICAM-1, VCAM-1, podoplanin (PDPN) and CD34^5^. Within the fibroblast clusters, we identified the presence of a population defined by the expression of *CD34, CCL19, TNFSF13B, ICAM1, VCAM1, CD82*, and *CXCL9* displaying key overlapping features consistent with the phenotype of immunofibroblasts^5^. Next, we identified a cluster defined by the expression of *CD55, C1R* and *ACKR3*, a chemokine scavenger receptor likely involved in the establishment of chemokine gradients within the TLS complex microdomains^36^. The potential role of this cluster in the cross talk with neighboring cell types was supported by the expression of the complement receptor *CD55* and *C1R*. Within the remaining three fibroblast subclusters, we identified a *SERPINE2* expressing subcluster, displaying expression of *CLEC14A, F2R* and elevated transcripts for *DKK3* indicating a potential engagement of Wnt signaling pathways. We also identified a *FOSB CDKN1A* expressing subcluster, displaying evidence of activation of the AP-1 pathway with increased *FOSB, FOS, JUND*, and *JUN* expression. Finally, an *ABCC9* fibroblast subcluster was recognized, defined by the expression of *APOC1, APOE, PLA2G2A*, and *CFD*, genes involved in lipid metabolism and alternative complement activation^37^ (fig 2b and suppl fig 2b). We used slingshot trajectory analysis with UMAP reduction to order cells and infer relationships between these clusters (suppl. fig 2b). The immunofibroblast cluster showed the closest relationship to the fibroblast SERPINE2 F2R and fibroblast FOSB CDKN1A clusters. The fibroblast ACKR3 cluster occupied one end of the trajectory and displayed similarities to the PI16 universal reservoir fibroblast population identified by Buechler *et al*^21^ including expression of *PI16, ACKR3, ACKR1, MFAP5, PCOLCE2, CD248*, and *IGFBP6*. Furthermore, the ACKR3 cluster displayed signatures associated with “stemness”, suggesting that the ACKR3 cluster might act as precursor of all the other fibroblast clusters. The ABCC9 and FOSB populations are the least distinct in terms of marker profile. They may represent activation or intermediate states between the other three populations, suggesting their nature as transitional/intermediate populations.

An additional stromal cell compartment was identified, characterized by the expression of *ACTA2* and therefore associated with mural cells or pericytes. Within this compartment, we classified 4 clusters of cells which comprised of 2 pericyte clusters and 2 vascular smooth muscle cell clusters (Fig 2a-c). Both pericyte clusters expressed the classical markers *RGS5, ACTA2*, and *MCAM*; however, one was further characterized by the expression of the lymphoid chemokines *CCL21* and *CCL19* and the glycoprotein tenascin C (*TNC*) (fig 2b and c). In contrast to other pericyte clusters, this CCL21^++^ CCL19^++^ TNC^+^ cluster expressed *CCL8, VCAM1*, and elevated *CCL2* gene transcripts (fig 2b and suppl. fig 2a). Whilst the CCL21^++^ CCL19^++^ TNC^+^ pericyte cluster expressed key chemokines implicated in TLS formation, it lacked expression of *TNFSF13B, CD82* and *PDGFRA* classical of immunofibroblasts. We named this population “immunopericytes”, to reflect the similarity to immunofibroblasts and the inferred immunological function of this population.

We extrapolated gene expression cassettes from single cell profiling and used it to validate the presence of the fibroblast and pericyte subclusters in the tissue using multiplex immunofluorescence (fig 2d), RNA scope and flow cytometry on tissue samples and digested SjS SG biopsies (Fig 2e and f and suppl. fig 2). Histological analysis demonstrated enrichment of CCL21+ pericytes in proximity to peripheral node addressing (PNAd^+^) high endothelial venules, supporting a potential immunological role in recalling T cells within the TLS for this population (suppl. fig 2e). To evaluate the developmental relationship between the immunofibroblasts and immunopericytes and explore the possibility that immunopericytes develop into immunofibroblasts during TLS establishment, we applied a fibroblast progenitor module score across fibroblast and mural populations, using the features identified by Buechler *et al* ^21^. Immunopericytes were found not to express this module, suggesting that they are unlikely to represent a stem fibroblast population in SG TLS (suppl. fig 2c).

Finally, to further dissect the distinct role of immunofibroblasts and immunopericytes in TLS, we performed gene-set enrichment analysis to identify differentially expressed pathways in these two populations. The top five enriched gene-sets for each cluster are represented in figure 2g. The immunofibroblast cluster showed enrichment for genes regulating antigen presentation and response to interferon gamma, reflecting the proinflammatory nature of this population. The fibroblast SERPINE2 F2R cluster was enriched for genes responsible for response to growth factors, ossification, and collagen metabolism. Fibroblast ABCC9 displayed enrichment of pathways involved in protein targeting endoplasmic reticulum. Fibroblast FOS CDKN1A showed enrichment for response to steroids and apoptosis. Finally, the ACKR3 CD55 cluster showed enrichment in the regulation of angiogenesis and enrichment for complement activation pathways. Conventional pericytes display enrichment for pathways responsible for sodium excretion; however, the immunopericyte population showed enrichment in pathways including transmembrane transport and ion transport, but not limited to sodium excretion.

### Different signals underpin the production of CCL21 and CCL19 in immunofibroblasts compared to immunopericytes

Intriguingly, whilst sharing the ability to secrete T cell chemokines, immunopericyte gene-set profiling was significantly different from that of immunofibroblasts and NFkB gene-set enrichment was restricted solely to the immunofibroblast subcluster (fig 3a). Accordingly, expression of *RELB* and *NFKB2* was limited to the immunofibroblasts, confirming that only immunofibroblasts, but not immunopericytes, rely on the lymphotoxin β receptor (LTβR) signalling pathway to produce lymphoid chemokines (fig 3b). Immunopericytes, on the contrary, show no enrichment in the genes of the non-canonical NFKB signalling pathway, and expressed both TNFα and IFNγ associated genes, such as *IFNGR1, IFNGR2, TNFRSF1A, IRF1* and *SOCS1* (fig 3b). We have previously demonstrated that in *ltβr*^*-/-*^ mice, the production of the lymphoid chemokines CCL19 and CCL21 was abrogated in immunofibroblasts^5^. However, at the time of our studies, we noted that *Ccl21* expression was unexpectedly maintained in vascular/perivascular structures, suggesting that LTβR signaling is not required to support perivascular expression of this T cell chemokine^5, 38, 39^ (fig 3c and suppl. fig 3). We have now explained this observation, by quantifying the presence of immunopericytes and immunofibroblasts in *wild type (wt) and ltβ*^*r-/-*^ murine SG infected with adenovirus to induce TLS^40^ (Fig 3d). As expected from our previous findings, in the absence of LTβR (and engagement of the non-canonical NFKB pathway), the number of mature immunofibroblasts that expressed lymphoid chemokines significantly decreased (fig 3d). However, the number of immunopericytes was not affected by LTβR abrogation, suggesting that, also in mice, the ability of this population to secrete lymphoid chemokines is independent from LTβR engagement.

**Figure 3:**
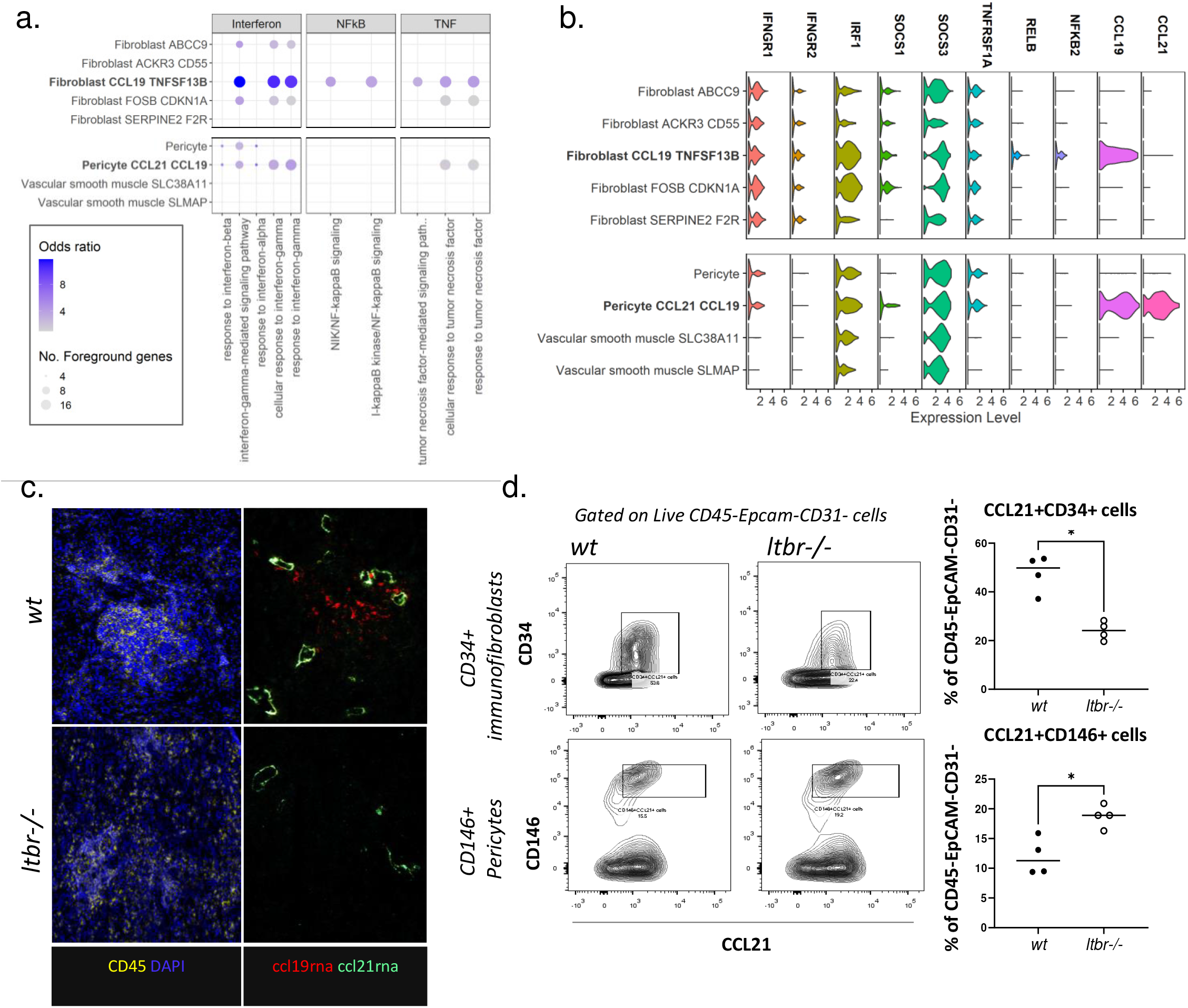
Immunofibroblasts and immunopericytes show differential dependence on signalling pathways: **a**, Select pathways from the geneset enrichment analysis (using GO and Kegg gene sets) of fibroblast and mural populations highlighting the enrichment of interferon and TNF pathways in both immunofibroblasts and immunopericytes, and NFκB pathway enrichment restricted to immunofibroblasts. **b**, Violin plot of expression of genes related to the enriched pathways shown in fig3a across fibroblast and mural clusters. **c**, Multiplex immunofluorescence image of ccl21rna (green), ccl19rna (red) vascular expression in DAPI (blue) and CD45^+^ (yellow) immune aggregate in TLS-induced salivary glands of *wt* and *Ltβr-/-* mice. *Ltbr-/-* results in abrogation of TLS and decreased CCL19 expression within immune aggregates. Perivascular expression of CCL19 remained in the *Ltβr-/-* **d**, Flow cytometric analysis of CCL21+CD34+ immunofibroblast and CCL21^+^CD146^+^ immunopericytes to confirm the dependence of immunofibroblasts on LTβR signalling and immunopericyte independence. Data are mean ±s.e.m from two independent experiments with two mice analyzed per group. \**p*< 0.05, unpaired *t* test.

### Single cell analysis defines cellular and functional properties for SjS and Sicca syndrome

SjS patients share a series of symptoms with patients affected by Sicca syndrome, a SG disease characterised by dryness in absence of the hallmark antibody profile and aberrant B cell activation typical of SjS. Importantly, whilst often presenting a diffuse T cell infiltrate in the SG, patients with Sicca syndrome do not form TLS^41^. We used single cell analysis applied to 6 Sicca syndrome SG and compared their profile to the SjS dataset (n=7) to dissect key cellular and functional properties of the two diseases. Pseudo-bulk analysis of the single cell data identified 1187 genes upregulated in SjS and 1016 genes upregulated in Sicca (fig 4a). Genes increased in SjS included *CXCL13, CXCR5, CD22, CD19, IGHG1, IL21, IFNG* and *SMR3B*. Genes upregulated in Sicca included *SCGB1D2, CAMP, SCGB2A1, S100A2*, and *LGALS7B* (fig 4a). In SjS, but not in Sicca samples, we identified a distinct *CCR6*^+^*LTB*^+^*BAFF*^+^ T helper cell cluster, a *CXCL13*^+^*CD40LG*^+^ T cell cluster and several B cell clusters (fig 4b and suppl. fig 4). Limited expression of *CD40* and *CD40L* was observed on all cell types in Sicca; *CD40*, on the contrary, was broadly expressed on SjS cells, underlining the relevance this pathway in SjS pathology^42, 43^.

**Figure 4:**
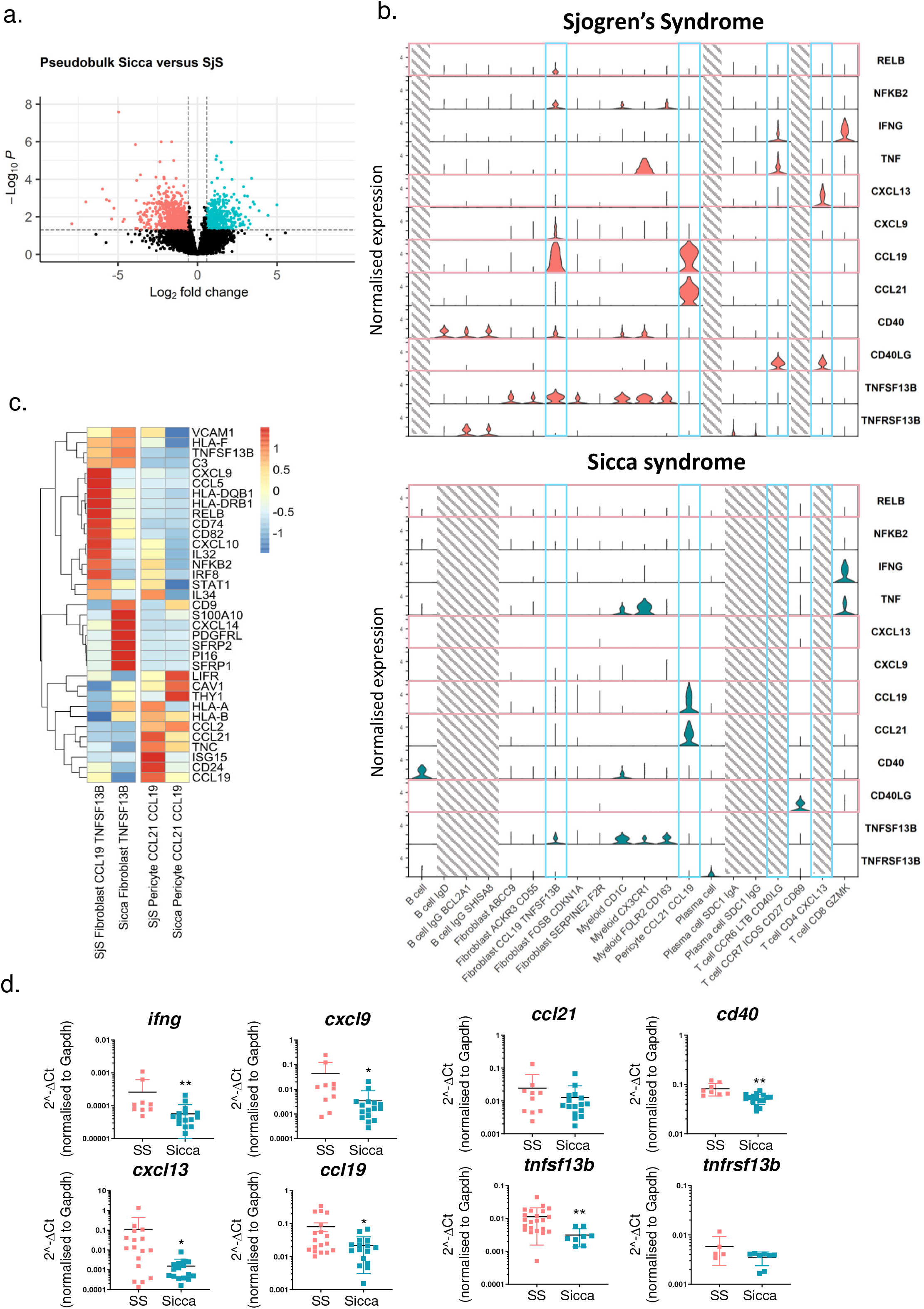
Signalling pathways differentiate between key populations in Sjogren’s syndrome and Sicca syndrome. **a**, Volcano plot illustrating differentially expressed genes detected between pseudobulked SjS and Sicca samples. **b**, Expression of genes related to signalling pathways and tertiary lymphoid structure formation in subsets from Sjogren’s syndrome or Sicca syndrome. **c**, Select differentially expressed genes detected between single-cell clusters of immunofibroblasts or immunopericytes between SjS and sicca, data are plotted as z-scores of the summed aggregate of all samples for each cluster including all samples. **d**, qPCR analysis for mRNA transcripts of *ifng, cxcl9, ccl21, cd40, cxcl13, ccl19, tnfsf13b and tnfrsf13b* in in whole tissue RNA extracts from SjS and Sicca biopsies. mRNA transcripts were normalized to the housekeeping gene *Gapdh* RNA transcripts. \**p*< 0.05; \*\**p*< 0.01, Mann–Whitney test. Data are representative as mean ±s.d., n=10-15.

Importantly, whilst immunopericytes were found in Sicca (though with lower expression of *CCL19*), classical immunofibroblasts were not detected in this condition. A population of fibroblasts expressing *TNFSF13B* but negative for the expression of the T cell attracting chemokines and *CD40* was present (fig 4b). Differential expression analysis among the two diseases (fig 4c) confirmed the diminished expression of the classical immunofibroblast signature in the Sicca *TNFSF13B* fibroblast cluster, increased expression of WNT-signalling related genes and overall, a different engagement of key genes defining the function of these cells.

Bulk mRNA analysis from whole tissue extracted from SG of patients affected by either SjS or Sicca confirmed that *IFNG, CXCL9, CXCL13, CCL19, CD40*, and *TNFSF13B* were all expressed at significantly higher levels in SjS compared to Sicca (fig 4d). In accordance, key differences in receptor/ligand interaction for both CXC and CC chemokines between in SjS and Sicca were identified and summarized in suppl. fig 4. Disease specific molecular properties in the two conditions associate with diverse fibroblast cluster distribution and, potentially, differentiation paths, establishing distinct molecular and cellular pathogenetic signatures for the two diseases.

### Distinct molecular properties identify GC harboring in secondary and tertiary lymphoid structures

TLS formation is a dynamic process. Small T cell aggregations occur first and are followed by progressive B cell infiltration and formation of germinal centers only occurs in mature, large TLS. It is known that lymphoid aggregates, commonly known in SjS as foci, at different stages of cellular organization are often found in the same SG tissue, limiting the ability of single cell analysis to resolve the cellular and molecular properties of TLS in the context of their spatial organization. To overcome this, we performed histology-guided laser microdissection of SjS SG foci defined by differing degrees of organization, using T/B cell area segregation and presence of fully formed CD21+ GC as key discriminators of evolving TLS^1^ (suppl. fig 5). Bulk sequencing and transcriptomic analysis were performed on isolated mRNA from non-segregated foci, segregated foci and foci with the formation of GC (TLS-GC). Tonsil germinal centers from healthy donors (tonsil-GC) were used as controls. PCA demonstrated sample clustering by level of maturation of the foci (Fig 5a and suppl. fig 5). To aid analysis we visualised expression of select genes known to be involved in lymphoneogenesis, co-stimulation, B cell apoptosis and survival (Fig 5b). Enrichment of genes involved in inflammatory pathways, inflammatory cell recruitment and co-stimulation was observed in foci displaying progressive degrees of organization. Interestingly, transcripts from organised foci and GC^+^ TLS overlapped with some of the key genes defining tonsil-GC. However, segregated foci and TLS-GC also presented a strong inflammatory signature with increased transcripts for *ICOS, CXCL10, INFG, TNF, CASP8* and proinflammatory chemokine receptors (fig 5b). As expected, classical tonsil GC genes included lymphotoxin, *CXCL13, AICDA* and regulators of apoptosis, known to be critical for GC homeostasis.

**Figure 5:**
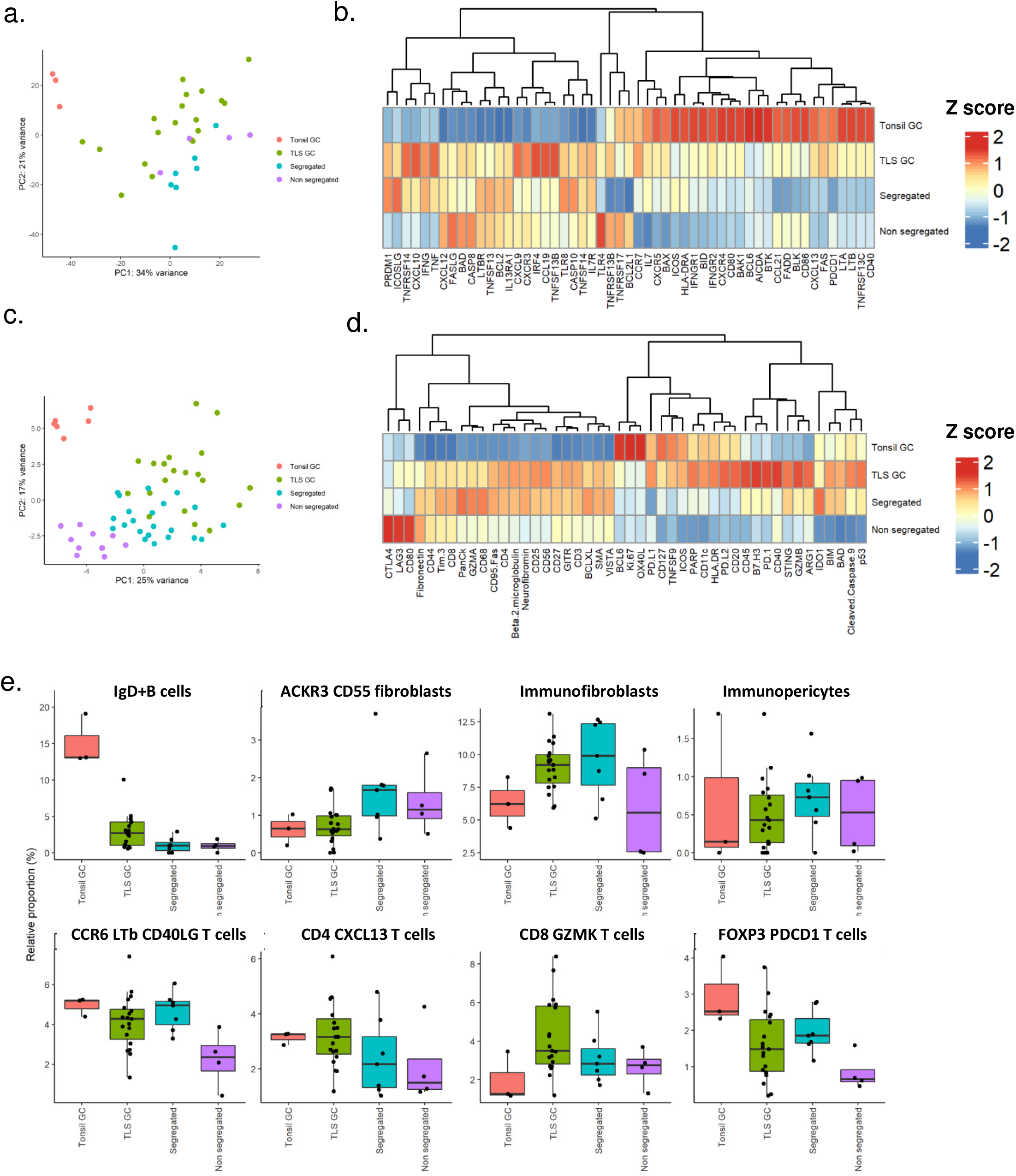
Bulk RNA-sequencing of microdissected regions of minor salivary glands from Sjogren’s syndrome patients and control tonsil tissue. **a**, PCA plot of bulk-sequencing of microdissection samples. **b**, Z-score mean expression of genes important to tertiary lymphoid organogenesis, in bulk-sequencing of microdissection data. **c**, PCA plot of Nanostring Geomx protein expression data. **d**, Z-score mean expression of proteins important to tertiary lymphoid organogenesis, in Nanostring Geomx data. **e**, Cibersortx inferred cell subset proportions in the bulk sequencing samples using the single-cell sequencing data as a reference atlas.

We confirmed these differences between classical SLO and TLS at the protein level using a Nanostring Geomx proteomic panel, the targets of which largely overlapped with the genes interrogated in fig 5b. Analysis was performed in regions of interest defined by the same criteria used to select areas for microdissection and bulk RNA-sequencing. PCA based on expression data of 49 proteins broadly recapitulated the clustering observed in figure 5a with tonsil-GCs clustering separately and the remaining samples showing a progression in clustering from small foci to segregated foci into mature TLS-GCs (Fig 5c). The heatmap in figure 5d illustrates clusters of proteins similar to those described for gene transcripts in figure 5b with CD127, OX40L, BCL, and Ki-67 forming a cluster with highest expression in Tonsil-GC. Protein expression for CD4, CD3, CD8, and CD68 displayed higher expression in all SG foci and TLS-GC than tonsil-GC along with higher expression of CD25, CD27, VISTA, and STING. Interestingly, higher expression of PD-1, PD-L1 and CD40 was detected in TLS-GC as compared to tonsil-GC (Fig 5d).

Bulk mRNA analysis on microdissected TLS provided key additional information to our single cell analysis. However, this approach lacks the cellular resolution necessary to attribute specific cell populations to histologically distinct, evolving TLS. We therefore used Cibersortx to generate a signature expression matrix and estimate cell subset abundance in the bulk RNA-sequencing samples (fig 5e, suppl. fig 5). As expected, estimated immunofibroblast cell proportions displayed a trend of increase from small foci to organized foci and TLS-GC samples. Interestingly, the potential precursors for this population (ACKR3^+^ cluster) appeared to be enriched in less mature TLS and progressively decrease with acquired organization and GC formation. Similarly, within the CD45^+^ leucocyte populations CD4 T follicular helper (T cell CD4 CXCL13) proportions progressively increased through the developmental stages of TLS with highest proportions in GC and tonsil samples. T helper 17-like (CCR6 LTB CD40LG T cells) estimated proportions followed a similar pattern, supporting the concept of a parallel development of both hematopoietic and stromal populations within TLS. To validate these data in tissue sections and define the spatially resolved cellular and structural components of TLS development, we performed hyperplex immunofluorescence imaging. Fully formed TLS and precursor (segregated & non-segregated) lymphoid organs were present throughout the tissue identifiable by prominent CD20^+^ B cell clusters and CD4^+^ T cell/CD20^+^ B cell interactions respectively (fig.6 a). We defined the developmental stage of each structure as T/B segregated with GC, T/B segregated or non-segregated Non-segregated structures showed variable B cell infiltration within a rich CD4^+^ T cell environment whereas segregated showed a rich B cell core with minimal CXCR5 expression and sparse CD4^+^ T cell infiltrate within the core but a rich T cell border. Full TLS formation showed a bed of CXCR5^+^ B cells, with CD21 expression and CXCL13^+^ cells with a spindle morphology, indicative of immunofibroblasts (fig. 6a). To quantify the differences in structure formation, the images were segmented to identify cell borders and the expression of each marker calculated for each cell. The expression profile of each cell identified 12 known phenotypes through single-cell clustering which were visualised on a UMAP and subsequently a spatial phenotype map (fig. 6b). Given the importance of structural organisation in these organs, we unbiasedly clustered tissue architectural features (cellular niches) using cellular neighbourhoods to define 5 neighbourhoods (as previously described). The enrichment of cells within these neighbourhoods was visualised on a heatmap (fig 6. d). Neighbourhood 1 was enriched for CXCR5^+^ B cells, TFH and immunofibroblasts representative of organised lymphoid structures. The abundance of neighbourhood 1 increased with increasing TLS developmental stage, further defining histological properties associated with full TLS maturation (fig 6. e and f).

**Figure 6:**
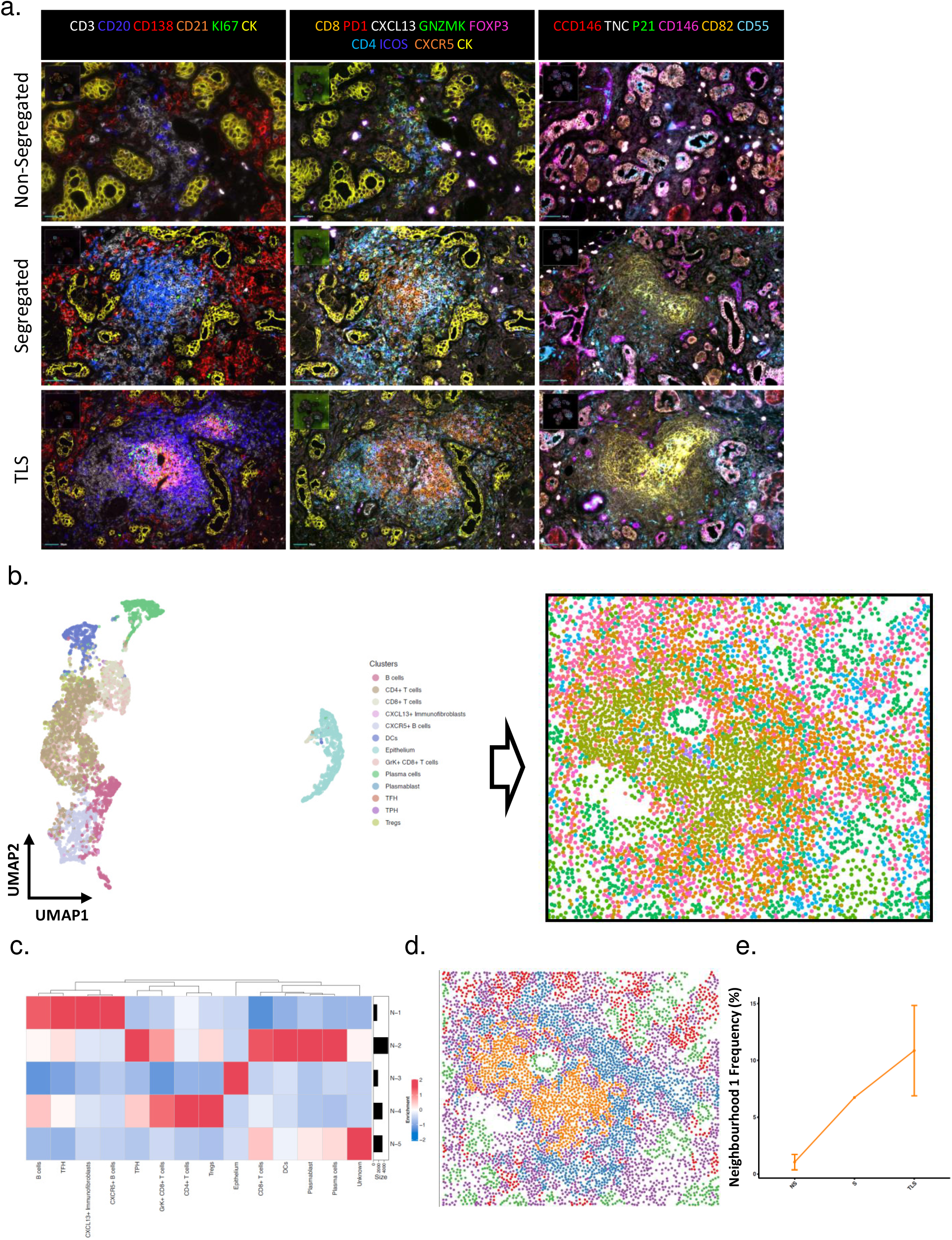
Single-cell spatial proteomics to identify spatially resolved cellular and structural landscape of TLS development. **a**. Multiplex immunofluorescence image of salivary gland highlighting the development of immune aggregates. **b**. UMAP of single-cell proteomic phenotyping and associated spatial phenotype map (of a TLS). **c**. Cellular neighbourhood heatmap **d**. Spatial neighbourhood map (of a TLS). **e**. Abundance of neighbourhood 1 (CXCR5+ B cell & Immunofibroblast enriched) as a function of developmental stage.

## Discussion

TLS are complex, evolving, multicellular structures able to shape the host immune response, impact disease trajectory and influence response to treatment in immune-mediated diseases and cancer^5, 31, 44, 45^. Previous reports have applied *in vitro* and *in vivo* approaches to characterize the contribution of various cellular populations to TLS formation. However, these studies, performed before the use of single cell analysis, did not exhaustively resolve TLS complex cellular composition, nor integrate these findings in the functional development of TLS^5, 11, 13^. In this study we have integrated single cell RNA sequencing, tissue transcriptomic and spatial proteomic data to deconvolute the cellular populations residing within human TLS, mapping spatial and functional interactions of TLS properties and their modulators as these structures evolve from small T cell aggregates to fully functional germinal centers.

We and others have previously identified fibroblasts as key elements for TLS establishment, as they precede and shape immune cell infiltration. In particular, we described a population of immunofibroblasts, stromal cells able to support lymphocyte migration and survival, while providing regulatory and structural function to the TLS. The origin of this population was, however, not demonstrated in humans. In our dataset we demonstrate the presence of a potential progenitor of TLS immunofibroblasts that is defined by the expression of *ACKR3* and *CD55* and shares key features with the PI16 universal fibroblast progenitor described by Buechler *et al*^21^. Interestingly, the transcripts encoded by the potential progenitor appeared to progressively decrease in increasingly organized TLS; whilst transcripts associated with more specialized fibroblasts increased. This phenomenon of fibroblast differentiation included acquisition of genes involved in the non-redundant functions of collagen metabolism and regulation of angiogenesis^14, 46^ and was confirmed in our data set and increased abundance of this population was detected in largely organized TLS and TLS harbouring functional germinal centers (GC+TLS). Importantly, the same population of immunofibroblasts was not identified in samples from patients with Sicca syndrome, a salivary gland disease, not characterized by TLS establishment. In our dataset we identified key differences in the properties specific to Sicca or SjS disease; in particular the enrichment in SjS, but not in Sicca, of specific cell populations (CXCL13 expressing T cells, LTB expressing T cells, and increased B cell and plasma cell state diversity)^12^. Moreover, we observed key variances in the differentiation of the fibroblast compartment, leading only in SjS and not in Sicca to the formation of mature immunofibroblasts. From our analysis it is not possible to establish whether these differences in fibroblast differentiation are due to differential imprinting in the common fibroblast progenitor or generated by the diverse microenvironmental cues present in the SG of patients with SjS and Sicca. In other autoimmune conditions, also characterized by TLS formation, production of autoantibodies has been reported to occur years prior to the establishment of tissue inflammation^12, 14^. The same autoantibodies have been involved in direct activation of non-immune cells^12^, suggesting the intriguing possibility that autoantibodies might condition fibroblasts to acquire a specific phenotype, able to support TLS formation, before the immune cells infiltrate the tissue. Animal data generated in rag^-/-^ mice appear to support this hypothesis, demonstrating early priming of immunofibroblasts in the absence of activated lymphocytes in tissue harboring TLS^5^.

Additional immunoregulatory properties could be ascribed to immunopericytes, a small population of mural cells that we identified in our dataset. Immunopericytes are differentiated from other pericytes by expression of the lymphoid chemokines *CCL21* and *CCL19*, two chemokines instrumental for naïve T cell recruitment and T cell zone organization in secondary lymphoid organs^11, 47, 48^. These immunopericytes do not express either LTβR or the genes classically associated with the alternative NFkB pathway^49^. Additionally, a population of perivascular reticular cells have previously been described with similarities to immunopericytes^50, 51^. In our dataset, progenitor potential in the immunopericytes was not demonstrated and, on the contrary, trajectory analysis appears to suggest a divergent differentiation pathway for immunofibroblasts and immunopericytes. Interestingly, immunopericyte presence precedes expansion of immunofibroblasts as Cibersortx analysis suggested that immunopericytes are present with similar abundance in small foci and organized or GC^+^ TLS. Moreover, immunopericytes are present in Sicca syndrome; however, in this condition, the cluster presents low expression of *CCL19*, most likely reflecting the decreased level of inflammation and lower transcripts of TNFα and IFNγ present in Sicca, that appears to support chemokine expression in this population. Given the totality of these data and the complexity of the cellular interactions unveiled by our bioinformatic analysis, we could speculate that, even in early inflammatory conditions, the differentiation of immunopericytes in close proximity to PNAd+ high endothelial venules, supports the formation of a functional unit for the migration of T lymphocytes, typical of small salivary gland foci. Persistence of inflammatory signals and progressive accumulation of lymphotoxin producing cells, in response to costimulatory signals and high TNFα and IFNγ production, results in further maturation of immunopericytes and progressive acquisition of immune regulatory properties by resident fibroblast progenitors. This process leads to the full differentiation of a novel population of immunofibroblasts, that, in turn become responsible for further recruitment of immune cells, including T follicular helper cells and B cells, providing not only spatial cues for cellular interaction, via chemokine production, but also survival factors and costimulation to incoming activated immune cells^11, 25, 35, 52^.

When first describing SLO formation, *Ansel et al*. reported the establishment of an amplificatory loop involving chemokines and lymphotoxin as a key element for the formation of the lymph node anlagen^53^. Here we demonstrate that the signals underpinning TLS development but also function are diverse. We expected the profile of highly organized TLS harboring GC to mimic the gene profile of classical SLOs. In contrast, transcriptomics and proteomic analysis demonstrated clear differences between ectopic and physiological lymphoid structures. Whilst some genes were shared, differences were observed in the expression of pro-survival and inflammatory genes, in genes regulating apoptosis, and in *AICDA*, the gene regulating somatic hypermutation and class switch recombination^54^. Taken together, these differences suggest that TLS are highly inflammatory hubs potentially characterized by inadequate mechanisms of control, as low AICDA has been associated with poor clone selection in the GCs. The absence of this signal, in association with abundance of survival factors (TNFSF13B, TNFSF14), inflammatory cytokines and chemokines (CXCL9, CXCL12) and costimulatory molecules (ICOS), unveiled by our analysis could explain the occurrence of malignant clone escape that has been described TLS and is responsible for the increased frequency in development of mucosal associated lymphoid tissue lymphoma observed in SjS.

Taken together, these data, provided us with important insights in the cellular and spatial complexity of TLS and the unique opportunity to compare the development of key TLS cellular elements in two diseases characterized by different pathogenic pathways in the same target organ. This has enabled us to define a trajectory of development of the non-hematopoietic, stromal cells during TLS development and to map the interactions of these populations with incoming lymphocytes in progressive stages of TLS assembly, allowing us to share the first comprehensive integrated analysis of autoimmune TLS development in humans. The importance of TLS in the context of non-autoimmune disease is just starting to be appreciated, as prognostic factors for disease progression and response to therapy, raising the need for a better understanding of signals and pathways involved in TLS establishment and maintenance in the tissue. We believe that the identification of TLS cellular and molecular properties and their relevant modulators provided by our analysis can be of importance for other diseases characterized by TLS formation and provide therapeutic cues for multiple conditions.

## Supplementary figure legends

**Supplementary figure 1: QC metrics of the SjS 10x data: a**, Expression of marker genes identifying gross-cell identities isolated from Sjogren’s minor salivary glands. **b**, Plots of the number of genes, number of UMIs, and percentage mitochondrial genes per sample. **c**, Proportions of gross cell states across samples

**Supplementary figure 2: Further characterisation of fibroblast and mural populations in SjS samples: a**, Expanded dotplot of previous and newly identified genes differentiating between fibroblast and mural clusters. **b**, Slingshot trajectory analysis of fibroblast clusters plotted against pseudotime or mapped to UMAP coordinates. **c**, Progenitor module score across fibroblast and mural populations generated using the Seurat function AddModuleScore with features identified as marking a progenitor population in the Buechler *et al* ^21^ dataset (specifically the genes *ACKR3, PI16, MFAP5, PCOLCE2, C3, PLA2G2A, IGFBP6, CD248*). **d**, CCL21 FMO control of flow cytometric identification of immunopericytes in human salivary glands. **e**, Multiplex immunofluorescence image identifying the *ccl21rna+ccl19rna+*CD146+ immunopericyte associates vessels within TLS in minor salivary gland tissue probed with a CD146 (grey), CD82 (yellow), PNAD (red) and DAPI (blue).

**Supplementary figure 3: a**, CCL21 FMO control of flow cytometric identification of immunopericytes in mouse salivary glands.

**Supplementary figure 4: Comparison of SjS and sicca single cell sequencing analysis. a** and **b**, UMAP reduction of SjS and sicca 10x data annotated with all cluster identities. **c**, Cellchat analysis chord diagrams showing sender and receiver populations for ligands included in CXCL or CCL pathways in both SjS and Sicca. Inner bars illustrate receiving populations and the proportion of the bar illustrates signal strength. **d, e**, and **f** show comparative cellchat analysis of immunofibroblasts (fibroblast TNFSF13B), immunopericytes (pericyte CCL21 CCL19), T cell, and B cell populations. T cell and B cell populations contained all subclusters of T cells and B cells in the granular analysis. **d**, circle plot of differential number of interactions in SjS versus Sicca. Red edges indicate increased number of interactions in SjS and blue in Sicca. Annotation provides the number of differential interactions. **e**, barplots providing the total number and strength of interactions in SjS versus Sicca. **f**, barplots of the relative number of interactions and of relative information flow in SjS versus Sicca by signalling pathways. Highlighted names indicate pathways with statistically significant differences.

**Supplementary figure 5: a**, CD3, CD20 and CD21 IHC staining showing representative images of non-segregated foci, segregated foci and TLS-GC selected from SjS salivary glands as well as Tonsil GC, for laser capture microdissection and subsequent RNA bulk sequencing. **b**, Extended plotting of CibersortX imputation of cell state proportions in bulk RNA-sequencing data derived from microdissection of SjS salivary gland tissue.

## Material and Methods

### Human samples

Labial minor salivary gland samples were obtained from patients recruited in the Optimising Assessment in Sjögren’s Syndrome (OASIS) cohort which recruits new patients attending the multidisciplinary Sjögren’s clinic at the Queen Elizabeth Hospital Birmingham, UK for assessment. Sjögren’s syndrome patients had a physician diagnosis of primary Sjögren’s syndrome and fulfilled the 2016 ACR/EULAR classification criteria. Participants with non-Sjögren’s sicca syndrome had signs and/or symptoms of dryness but did not have a physician diagnosis of SS or fulfill 2016 classification criteria. Salivary gland biopsy samples were divided in two: one for the scRNAseq study and the second for histological analysis to confirm diagnosis. Histological diagnosis is reported as presence of focal lymphocytic sialadenitis (FLS, suggestive of Primary Sjögren’s Syndrome, PSS) or non-specific chronic sialadenitis (NSCS), in the case of non-Sjögren’s sicca syndrome. All OASIS participants provided written informed consent and the study was approved by the Wales Research Ethics Committee 7 (WREC 7) formerly Dyfed Powys REC; 13/WA/0392.

### Mice and salivary gland cannulation

C57BL/6 wild-type *(wt)* mice were purchased from Charles River. *Ltbr*^*-/-*^ mice were provided by Jorge Caamano. All mice were maintained under specific pathogen-free conditions in the Biomedical Service Unit at the University of Birmingham according to Home Office and local ethics committee regulations. Under ketamine/domitor anaesthesia, the submandibular glands of female C57BL/6 and knock-out mice (8-12 weeks old) were intra-ductally cannulated with 10^8^-10^9^ p.f.u. of luciferase-encoding replication-defective adenovirus (Adv5) as previously described^5, 13, 40.^

### Isolation of cells from salivary glands

Minor salivary gland biopsies were taken surgically from the lip and frozen in 1mL of CryoStor® CS10 (StemCell Technologies) at −80°C. For preparation of single-cell suspension, firstly the frozen tissue sample in Cryotube were quickly thawed in water bath at 37°C and washed twice in pre-warmed 5%FBS RPMI media. The salivary gland biopsies were then enzymatically digested as previously described^5^. Dead cells were removed using the EasySepTM Dead Cell Removal (Annexin V) kit from the digested samples following manufacturer’s instructions before proceeding for the scRNA sequencing using the 10x platform.

### Single-cell RNA sequencing

Library preparation and sequencing was completed by Oxford Genomics Centre. Cell counts in each sample were quantified using the Bio-Rad TC20. 10,000 cells per sample were used to generate libraries with the 10x Genomics v3 Single Cell 3’ kit before sequencing to a minimum depth of 50,000 reads per cell using the NovaSeq 6000.

### Single-cell sequencing alignment and pre-processing

Reads for each sample were aligned to the hg38 transcriptome and quantified using Cellranger v3.0.2 on the University of Birmingham BlueBEAR High Performance Computing service. Potential doublets were identified and removed using Scrublet v0.2.1 prior to sample aggregation and UMI normalisation using scripts adapted from the Sansom lab repository (https://github.com/sansomlab). Cells with greater than 200 unique genes expressed of which less than 20% are mitochondrial genes were taken forward for analysis using R v3.6.1 and Seurat v.2.3.4 ^55^. Sjogren’s syndrome samples and Sicca syndrome samples were analysed independently from this point following parallel analysis pipelines. Data were log normalised, the effects of the number of UMIs and percentage mitochondrial genes were regressed out before scaling and centring the data. For the comparative analysis of Sjogren’s syndrome and Sicca syndrome single-cell datasets, seurat objects were merged and renormalised and scaled.

### Cluster identification

Variable genes were identified by selecting outliers on a mean variability plot (Seurat FindVariableFeatures). Principal component analysis was completed before using Harmony v1.0 to minimise the effect of donor on subsequent clustering. Dimensional reduction was completed using the UMAP algorithm based on the harmony alignment. Clustering using the Seurat function FindClusters was investigated across resolutions from 0.2-1.2 and stability was assessed using Clustree v0.4.3. After gross cell types were identified the data was subset to each cell type and the process was iterated to identify subclusters. Final clustering was completed in R v4.0.3 and Seurat v4.0.2.

### Single-cell sequencing ambient background correction

To account for background noise with the single-cell sequencing dataset we implemented the probabilistic archetype assignment method from^22^. Cells with a larger probability of ambient noise than any other cell archetypes were removed from the dataset. The ambient probability for each cell was used in models for differential expression analysis during subclustering iterations.

### Differential expression analysis

Differential expression between cell clusters was completed using the Seurat function FindClusters with the MAST algorithm using the ambient probability as a latent variable within the model.

### Single-cell sequencing trajectory analysis

R v4.0.3 and slingshot v1.8 were used to run trajectory analysis on the fibroblast clusters in UMAP space.

### Gene set enrichment

Gene-set enrichment analysis of the fibroblast and mural subsets was run using gsfisher v0.2 (https://github.com/sansomlab). Features were selected for each cluster using the FindAllFeatures function (Seurat v4.0.3) running MAST with the calculated ambient probability as a latent variable. Enrichment was tested against GO terms and Kegg pathways.

### Single-cell sequencing ligand-receptor analysis

CellChat v1.4 was used to run ligand-receptor analysis on the single-cell sequencing dataset.

### Multiplex immunohistochemistry staining protocol and image acquisition

3 μm thick sections of FFPE tissue on super frost plus slides were sectioned. Indirect detection by fluorescence was based on the Opal™ Multiplex IHC method (Akoya), performed on the Autostainer Leica Bond RX and Comet (Lunaphore). For staining in Comet, FFPE sectioned were deparaffinised and were subjected to a primary heat-induced epitope retrieval (HIER) in Tris EDTA buffer, pH 9.0 120°C for 60minutes in the PT-link (Lunaphore). Antibodies used were CD45, VE-cadherin, podoplanin, CD90, CD146, CD3, CD20, CD8, CD4, CD68, CD11c, PNAD, CD82, Tenascin C, pan-cytokeratin, CD138, CD21, GranzymeK, P21, CD55, FOXP3, ICOS, PD1, CXCL13, CCD146, CXCR5 and KI67. Immunofluorescent signal was visualized using the OPAL™ 7-color fIHC kit (Perkin Elmer, MA) TSA dyes 480, 520, 570, 620, 690 and 780 counterstained with Spectral DAPI. Or Goat anti-mouse IgG/IgM Alexa Fluor 555 and goat anti-rabbit Ig Alexa Fluor 647. Multispectral images were acquired at ×20 magnification using the Vectra Polaris Automated Quantitative Pathology Imaging System (Akoya) or COMET (Lunaphore). MoTIF settings were used for multispectral image acquisition. Multispectral image processing of multiplex IHC stains was performed using Phenochart (version 1.0.11/Akoya) and inForm Image Analysis Software (version 2.3, Akoya) or Lunaphore image viewer.

### COMET multiplex IHC Image Processing and Cellular Segmentation

After acquisition, images were shading corrected, cycle aligned, stitched and merged into OME-TIFF pyramidal format within the Lunaphore COMET Explorer software. Autofluorescence for TRITC and Cy5 was conducted in the Lunaphore COMET Viewer v2. Single channels from each OME-TIFF stack were then extracted and regions containing lymphoid structures were sampled and saved as individual channel tiff files. Each region was segmented with CellSeg using default settings ^56^, to identify each cell nucleus and membrane, and the segmentation mask for each was saved. These masks were then used to extract the expression, spatial and morphological information of each cell for the individual channel images. For nuclear channels (e.g., DAPI, Ki67 & FoxP3), the nuclear mask was used to extract cellular information. This information was saved as an FCS-like tabular data file.

### Cell Phenotype Identification and Neighbourhood Analysis

FCS-like files generated from COMET imaging were imported into R. Expression values were compressed to the 99.9th percentile per region. The morphology metrics cell size, solidity and eccentricity were z-scored and used as initial quality control steps to remove segmentation artefacts. Cell phenotyping was performed using CELESTA on each with a custom generated cell expression matrix ^57 57^. The phenotype assignments were qualitatively validated by plotting them in their original spatial location and comparing the patterns to the images. The phenotypes were then visualised on a UMAP. Cells were assigned positive or negative for each functional protein by the findCutoff function of MetaCyto. Cellular neighbourhoods were calculated by recording the 10 nearest neighbours of each cell and clustering these using k-Nearest Neighbours (kNN) with k = 6 as previously described ^58^.

### RNAscope

Indirect detection by fluorescence was based on the Opal™ Multiplex IHC method (Akoya) and performed on the Autostainer Leica Bond RX. Samples were probed for RNAscope^®^ LS 2.5 Probe - Hs-CCL19, Hs-CCL21, Ms-CCL19 and Ms-CCL21 transcript variant 1 mRNA following manufacturer’s instructions (ACDBio). Subsequent staining was performed with antihuman CD146 (Abcam) CD45(Cell Signalling) antibodies. Staining was performed using the Opal™ 7-Color Automatic IHC Kit (Akoya) according to the manufacturer’s recommendations. Signal amplification and covalent binding of fluorophore was achieved by using a tyramide signaling amplification reagent (included in the Opal kit) that is conjugated with a different fluorophore for each cycle. Multispectral images were acquired at ×20 magnification using the Vectra Polaris Automated Quantitative Pathology Imaging System (Akoya). MoTIF settings were used for multispectral image acquisition. Multispectral image processing of multiplex IHC stains was performed using Phenochart (version 1.0.11/Akoya) and inForm Image Analysis Software (version 2.3, Akoya).

### Microdissection and Bulk-RNA sequencing

The selected tissue samples for microdissection processing were acquired from patients recruited within the OASIS cohort at the University of Birmingham, under ethics number 10-018. Preparation of the samples for microdissection: all surfaces of the cryostat and lab bench were thoroughly cleaned with 70% Etoh and sterilised with ultraviolet (UV) light for at least 30 min. Containers used to keep store the collected samples and the filter paper used to separate slides (Membrane slides, Carl Zeiss™ 1.0 PEN NF 41590-9081-000) must also be treated with UV for 30 min and kept sterile before handling the tissue. To cut tissue for microdissection, the UV sterilised membrane slides were maintained at a cool temperature within the cryostat prior to sample cutting. The tissue was mounted on a cryostat block with as little mounting media (OCT, TissueTeK) as possible to minimise interference with laser cutter during microdissection. Sections were cut at 8 um thickness and transferred to the membrane slides. Once the section had been mounted onto the membrane slides, the membrane slide was stored in its carrier box and in dry ice until final storage at -80°C prior to staining with Cresyl violet acetate (SIGMA). Preparation of Cresyl violet acetate is required one day prior to staining the mounted tissues. Preparation of the CresylViolet (1% solution) included dissolving the powder in 80% ethanol which had been made using sterile RNase-free water in a MSC Class II ventilator system. This solution was mixed overnight on a shaker at 4°C to ensure that the CresylViolet was completely dissolved, and the solution was then passed through a 70μm strainer before use. The slides were first dipped in RNase-free water and suspended for 2 min in 70% Etoh, also prepared in RNase-free water, to facilitate the removal of OCT on the tissue. The slides were then suspended in the CresylViolet 1% solution for 30 secs to 1 min and excess solution was tapped off. The slides were then processed for 1 min in 100% Etoh and this step was repeated three times. The PALM slides were then left to dry until the membrane was observed to be completely dry. The PALM slides were then stored at -80°C in a parafilm sealed 50 ml falcon until the samples were ready for microdissection.

Prior to microdissection, the PALM slides were bought to room temperature for 10 min. Microdissection was completed using the PALM Robo Software V.4.6 software using the brightfield “AxioCam CC1” setting. The objective lens used to micro dissects SG sections was 10X whilst tonsil was set at 5X. The area of interest was then identified using the ocular eye piece. Once the region had been selected the laser was aligned to the section that was to be cut. The collection tubes required for the capture of the microdissected tissue contained 20 ul of RLT buffer with B-mercaptanol, ensuring the lid was kept dry, and tube was kept on dry ice prior to use. The collection tubes were then mounted onto the *Collector Set 200 CMII* and set to the correct position to collect the microdissected sample as they were cut by applying the calibration procedure to collect tissue in the centre of the lid of the collection tube which were stored at -80°C prior to RNA extraction. RNA extraction from the microdissected samples was completed as per the protocols provided using the RNeasy FFPE kit (Qiagen).

Library preparation and sequencing from extracted RNA was completed by Genomics Birmingham. Initial libraries were prepared using the SMART-Seq v4 Ultra Low Input RNA Kit. Further samples were processed using the NEBNext rRNA Depletion Kit v2 and NEBNext Ultra II Directional RNA Library Prep Kit for Illumina with NEBNext Multiplex Oligos for Illumina. All libraries were sequenced using a NextSeq 500 sequencer. Adapters were trimmed with trimmomatic v0.39, reads aligned to the hg38 transcriptome with STAR v2.7.2b, and counts summed using featureCounts in Subread v2.0.1 using the University of Birmingham BlueBEAR High Performance Computing service. R v4.0.3 and DESeq2 v1.30.1 were used for downstream analysis. To minimize batch effects during differential expression analysis batch was included as a factor in the DESeq2 model and for principal component analysis and visualisations batch effects were minimised in variance stabilising transformed data using the limma v3.46.0 function removeBatchEffect.

### Geomx Nanostring

The Nanostring GeoMX Digital Spatial Profiler (DSP) Protein slide preparation was followed according to the manufacturer’s instruction (https://nanostring.com/products/geomx-digital-spatial-profiler/geomx-protein-assays). Briefly, the slides were baked, deparaffinized, and antigen retrieval was performed using Citrate buffer pH 6.0. Next, the samples were incubated with 84 oligo-labeled primary antibodies: a Human Immune Cell Profiling Core, a Human Immune Activation Status Panel, a Human Immune Cell Typing Panel, a Human Immune Cell Death Panel and a Human Immune Drug IO Panel. The same sections were then stained with fluorescently labeled SYTO13, CD45, CD21 and CD138 to identify tissue regions of interest (ROI). The slides were then loaded onto the GeoMX DSP and 4-6 ROIs were randomly selected per slide and analyzed for protein expression. All the indexing oligonucleotides were collected into a 96 well plate and were then hybridized to fluorescent barcodes using GeoMX Hyb Codes. After hybridization, samples were processed using the using the nCounter system according to the manufacturer’s instructions. 49 regions of interest passed QC criteria and were taken forward for analysis. Using the Geomx software we scaled the data to number of nuclei/ROI and subsequently normalised using negative control probes. Data were exported and PCA analysis and heatmaps were generated using R v4.0.3 and the pheatmap v1.0.12 package.

### Cell type abundance estimation

Using the deconvolution tool Cibersortx ^59^, the Sjogren’s syndrome single-cell data was used as a reference dataset to generate a signature matrix which was used to impute cell type abundance in the bulk RNA-sequencing samples. 100 permutations were run for the imputation.

## Acknowledgments

We thank Jorge Caamano for providing animals; Technology Hub (University of Birmingham) for flow cytometry, imaging, and PCR, Birmingham Environment for Academic Research (University of Birmingham) for providing HPC infrastructure, Genomics Birmingham (University of Birmingham) and Oxford Genomics Centre (University of Oxford) for sequencing services, and Birmingham Tissue Analytics (University of Birmingham) for assistance with RNAscope analysis, Geomx support and advanced IHC multiplexing platforms (Comet and Vectra). We are indebted to the Biomedical Services Unit and Biological Services Facility for maintaining the colonies. F.B. was funded by the Wellcome Trust, Arthritis Research UK (ARUK) Senior Fellow and NECESSITY Consortium. C.D.B. was supported by ARUK Grant (G0601156); B.A.F., S.J.B and SN have received support from the National Institute for Health Research (NIHR) Birmingham Biomedical Research Centre at the University Hospitals Birmingham NHS Foundation Trust and the University of Birmingham (Grant BRC-1215-20009) and patients were recruited at the NIHR/Wellcome Trust Birmingham Clinical Research Facility. The views expressed are those of the authors and not necessarily those of the NIHR or the Department of Health and Social Care. SN has received support from Wellcome Trust Fund for the Geomx assay. APC is supported by a Kennedy Trust for Rheumatology Research Senior Clinical Fellowship.

