## Supplementary Table S1 for "A cellular and spatial map of salivary glands at single cell resolution reveals the functional basis of tertiary lymphoid structure formation in Sjogren’s syndrome"

| SampleID | Sex | Diagnosis | Presence_of_antiRo_antibodies | Focus_score | Histology |
|---|---|---|---|---|---|
| SalivaryGland1 | F | PSS | Neg | 1.74 | FLS. Mild focal fibrosis. |
| SalivaryGland2 | M | PSS | Neg | 2.78 | FLS. Mild focal fibrosis. |
| SalivaryGland3 | F | sicca | Neg | 0 | NSCS. Small aggregate and dispersed population of T and B cells. Diffuse fibrosis. |
| SalivaryGland4 | F | PSS | Neg | 1.29 | FLS. Mild focal fibrosis. |
| SalivaryGland5 | F | PSS | Neg | 2.6 | FLS. Mild focal fibrosis. |
| SalivaryGland6 | F | PSS | Pos | 1.32 | FLS. Mild focal fibrosis. |
| SalivaryGland7 | F | sicca | Neg | 1.3 | NSCS. Diffuse marked fibrosis. |
| SalivaryGland8 | M | sicca | Neg | 0 | Near normal, few sparse T cells |
| SalivaryGland9 | F | sicca | Neg | 0 | Few small aggregates on immunohistochemistry. Focal marked fibrosis. |
| SalivaryGland10 | F | sicca | Neg | 0.74 | NSCS. One aggregate on H&E, diffuse CD3 and small aggregates on immunohistochemistry. Focal marked fibrosis. |
| SalivaryGland11 | M | sicca | Neg | 0 | Minor inflammatory changes only, mild focal fibrosis. |
| SalivaryGland12 | F | PSS | Pos | 0.2 | Mild sialadenitis on H+E. Good numbers of T cells on immunohistochemistry, diffusely distributed and forming small aggregates. Mild focal fibrosis. |
| SalivaryGland13 | F | sicca | Neg | 0.81 | Mild sialadenitis. Minimal focal fibrosis. |
| SalivaryGland14 | F | PSS | Pos | 2.2 | FLS |
