## Supplementary Figures for "A cellular and spatial map of salivary glands at single cell resolution reveals the functional basis of tertiary lymphoid structure formation in Sjogren’s syndrome"

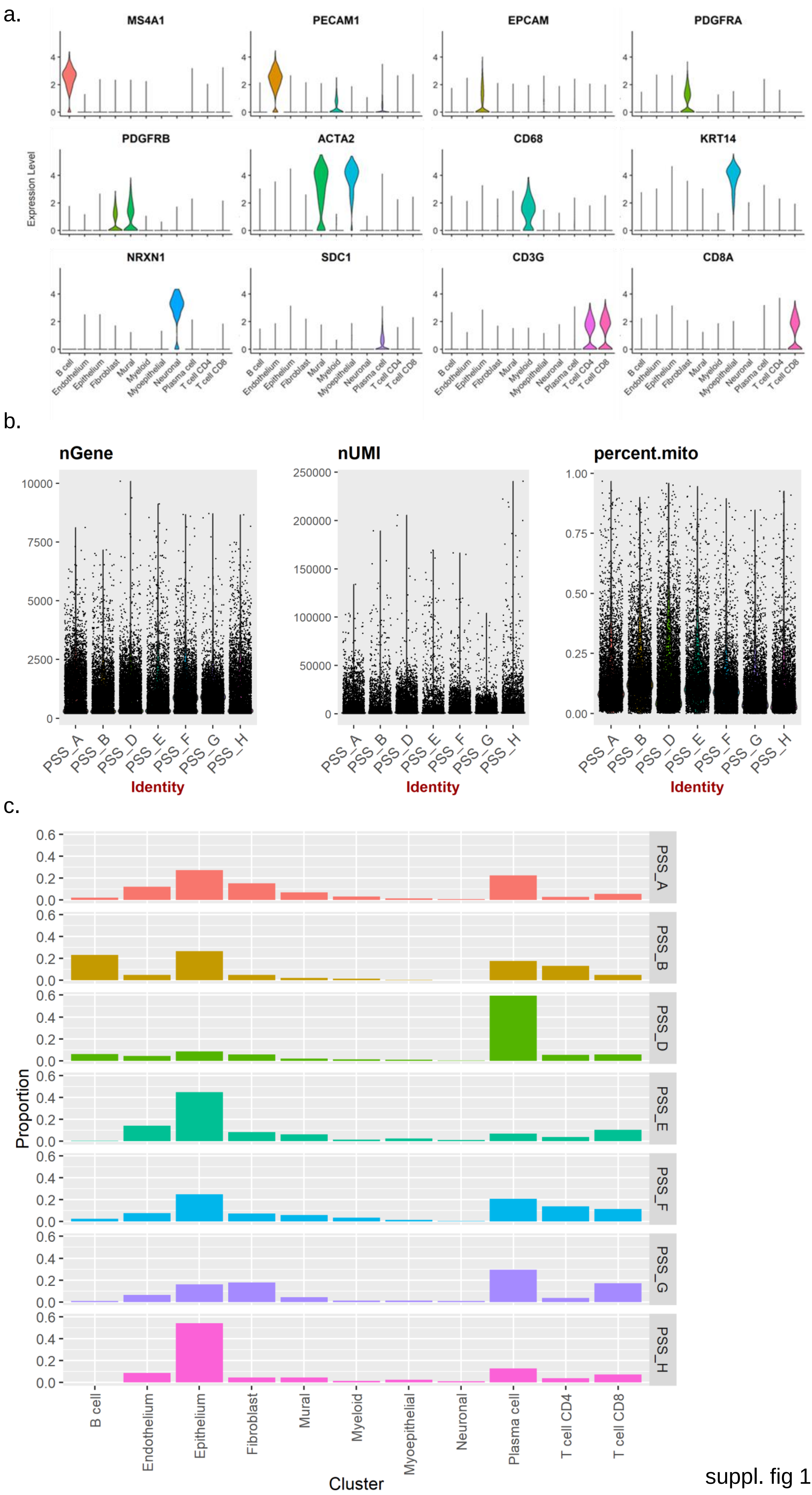
a.
MS4A1
PECAM1
EPCAM
PDGFRA
PDGFRB
ACTA2
CD68
KRT14
NRXN1
SDC1
CD3G
CD8A
Expression Level
B cell
Endothelium
Epithelium
Fibroblast
Mural
Myeloid
Myoepithelial
Neuronal
Plasma cell
T cell CD4
T cell CD8
b.
nGene
nUMI
percent.mito
PSS_A
PSS_B
PSS_D
PSS_E
PSS_F
PSS_G
PSS_H
Identity
c.
Proportion
Cluster

suppl. fig 1

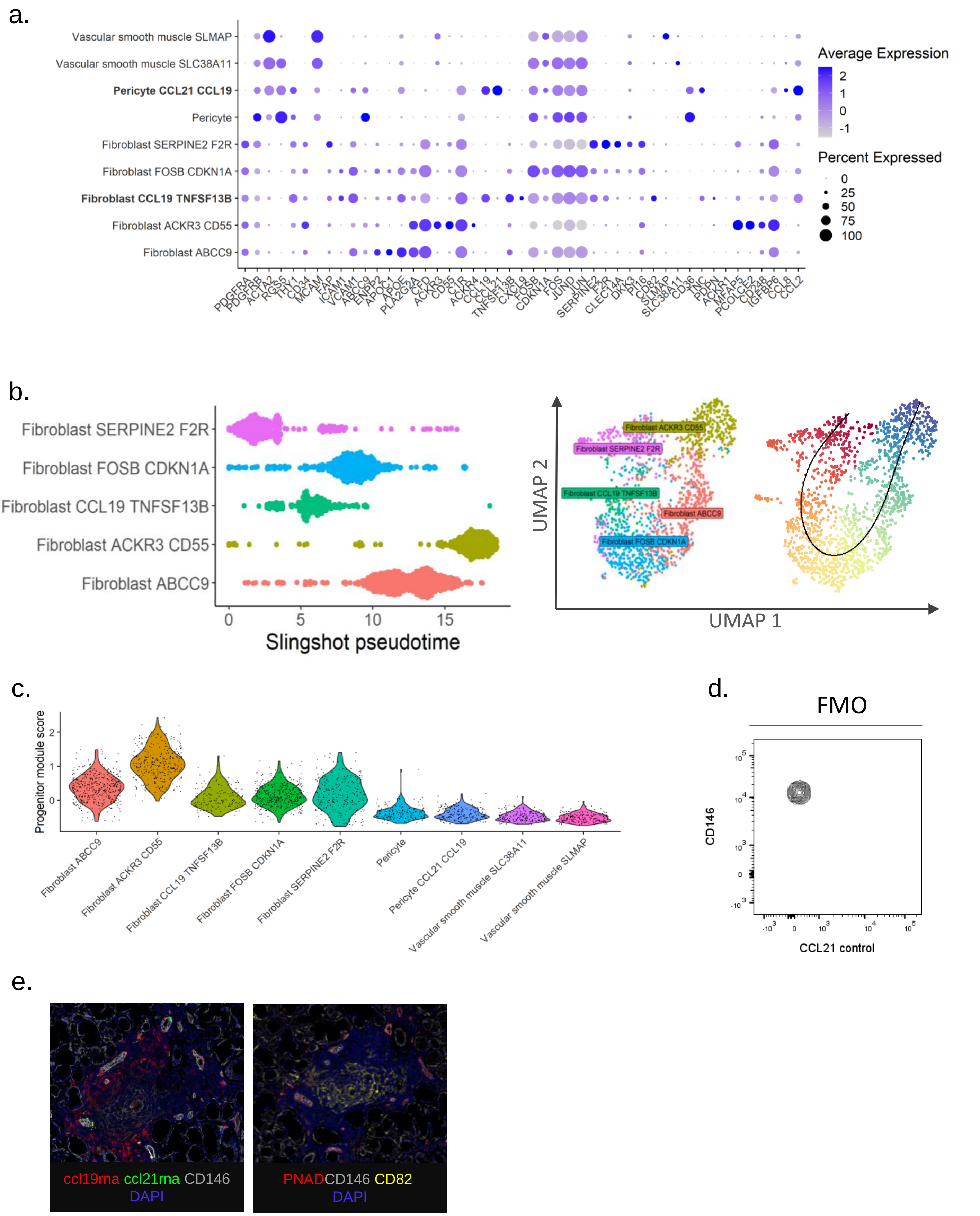
a.
Vascular smooth muscle SLMAP
Vascular smooth muscle SLC38A11
Pericyte CCL21 CCL19
Pericyte
Fibroblast SERPINE2 F2R
Fibroblast FOSB CDKN1A
Fibroblast CCL19 TNFSF13B
Fibroblast ACKR3 CD55
Fibroblast ABCC9
PDGFRA PDGFRB ACTA2 RGS5 THY1 CD34 MCAM FAP ICAM1 VCAM1 ABCC9 ENPP2 APOC1 APOE PLA2G2A C7 CFD ACKR3 CD55 C1R ACKR4 CCL19 CCL21 TNFSF13B CXCL9 FOSB CDKN1A FOS JUND JUN SERPINE2 F2R CLEC14A DKK3 PI16 CD82 SLMAP SLC38A11 CD36 TNC PDPN ACKR1 MFAP5 PCOLCE2 CD248 IGFBP6 CCL8 CCL2
Average Expression
2
1
0
-1
Percent Expressed
0
25
50
75
100
b.
Fibroblast SERPINE2 F2R
Fibroblast FOSB CDKN1A
Fibroblast CCL19 TNFSF13B
Fibroblast ACKR3 CD55
Fibroblast ABCC9
0
5
10
15
Slingshot pseudotime
UMAP 2
UMAP 1
Fibroblast ACKR3 CD55
Fibroblast SERPINE2 F2R
Fibroblast CCL19 TNFSF13B
Fibroblast ABCC9
Fibroblast FOSB CDKN1A
c.
Progenitor module score
2
1
0
Fibroblast ABCC9
Fibroblast ACKR3 CD55
Fibroblast CCL19 TNFSF13B
Fibroblast FOSB CDKN1A
Fibroblast SERPINE2 F2R
Pericyte
Pericyte CCL21 CCL19
Vascular smooth muscle SLC38A11
Vascular smooth muscle SLMAP
d.
FMO
CD146
CCL21 control
e.
ccl19rna ccl21rna CD146
DAPI
PNAD CD146 CD82
DAPI

suppl. fig 2

a.

FMO

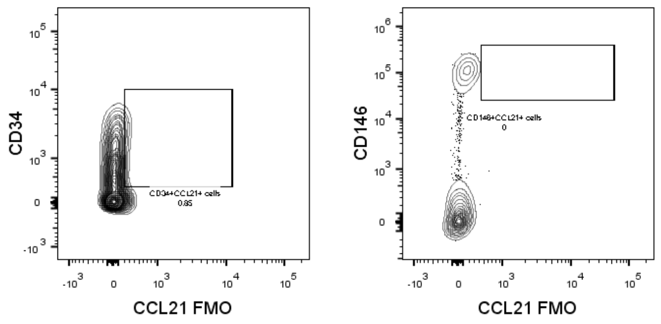
CD34
CD34+CCL21+ cells
0.85
CCL21 FMO
CD146
CD146+CCL21+ cells
0
CCL21 FMO

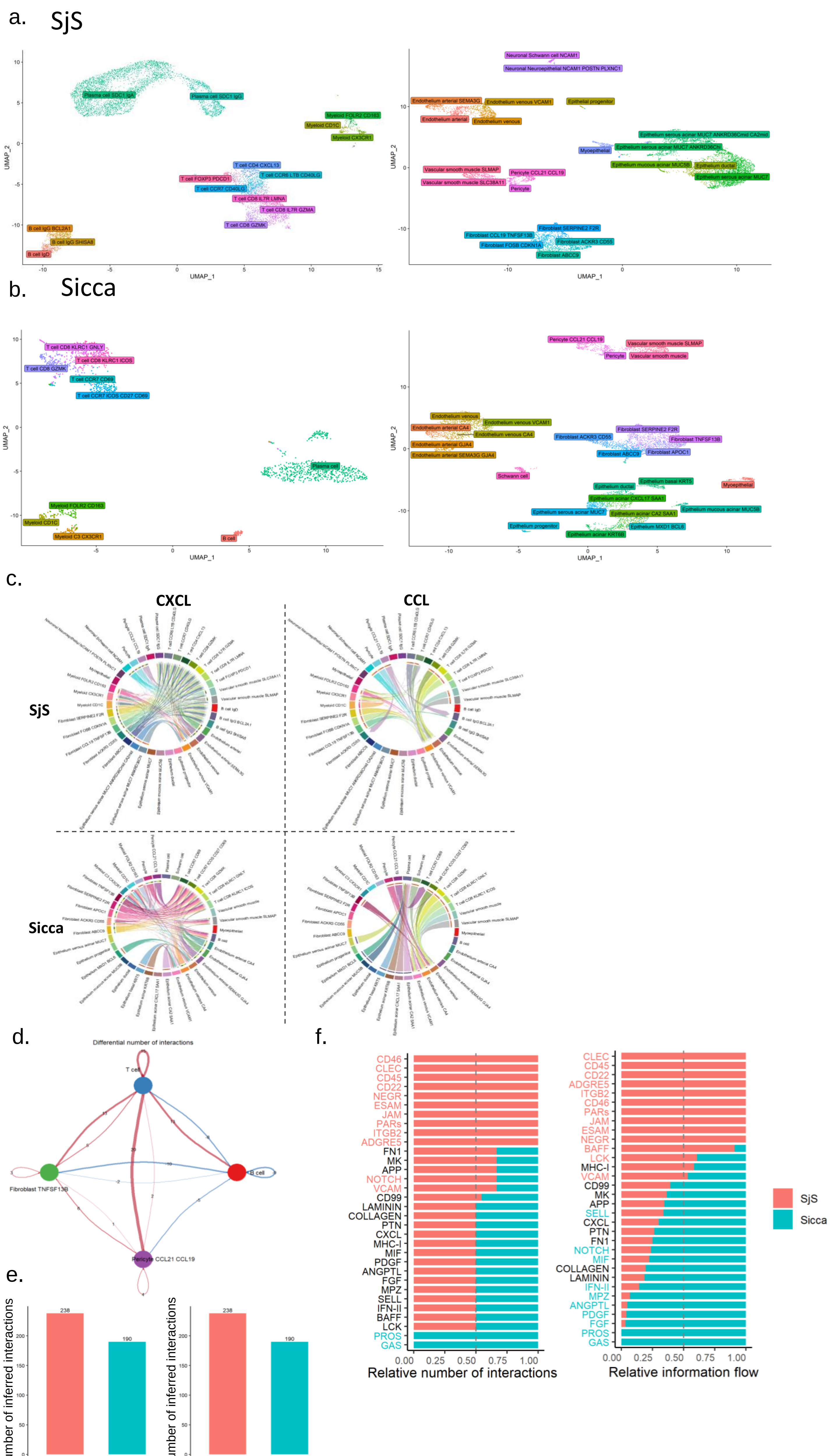
a. SjS
Plasma cell SDC1 IgA
Plasma cell SDC1 IgG
Myeloid FOLR2 CD163
Myeloid CD1C
Myeloid CX3CR1
T cell CD4 CXCL13
T cell CCR6 LTB CD40LG
T cell FOXP3 PDCD1
T cell CCR7 CD40LG
T cell CD8 IL7R LMNA
T cell CD8 IL7R GZMA
T cell CD8 GZMK
B cell IgG BCL2A1
B cell IgG SHISA8
B cell IgD
Neuronal Schwann cell NCAM1
Neuronal Neuroepithelial NCAM1 POSTN PLXNC1
Endothelium arterial SEMA3G
Endothelium venous VCAM1
Epithelial progenitor
Endothelium arterial
Endothelium venous
Epithelium serous acinar MUC7 ANKRD36Cmid CA2mid
Myoepithelial
Epithelium serous acinar MUC7 ANKRD36Chi
Epithelium mucous acinar MUC5B
Epithelium ductal
Epithelium serous acinar MUC7
Vascular smooth muscle SLMAP
Pericyte CCL21 CCL19
Vascular smooth muscle SLC38A11
Pericyte
Fibroblast SERPINE2 F2R
Fibroblast CCL19 TNFSF13B
Fibroblast ACKR3 CD55
Fibroblast FOSB CDKN1A
Fibroblast ABCC9
UMAP_1
UMAP_2
b. Sicca
T cell CD8 KLRC1 GNLY
T cell CD8 KLRC1 ICOS
T cell CD8 GZMK
T cell CCR7 CD69
T cell CCR7 ICOS CD27 CD69
Plasma cell
Myeloid FOLR2 CD163
Myeloid CD1C
Myeloid C3 CX3CR1
B cell
Pericyte CCL21 CCL19
Vascular smooth muscle SLMAP
Pericyte
Vascular smooth muscle
Endothelium venous
Endothelium venous VCAM1
Endothelium arterial CA4
Endothelium venous CA4
Endothelium arterial GJA4
Endothelium arterial SEMA3G GJA4
Fibroblast SERPINE2 F2R
Fibroblast ACKR3 CD55
Fibroblast TNFSF13B
Fibroblast ABCC9
Fibroblast APOC1
Schwann cell
Epithelium basal KRT5
Epithelium ductal
Myoepithelial
Epithelium acinar CXCL17 SAA1
Epithelium serous acinar MUC7
Epithelium acinar CA2 SAA1
Epithelium mucous acinar MUC5B
Epithelium progenitor
Epithelium MXD1 BCL6
Epithelium acinar KRT6B
UMAP_1
UMAP_2
c.
CXCL
CCL
SjS
Sicca
d.
Differential number of interactions
T cell
Fibroblast TNFSF13B
B cell
Pericyte CCL21 CCL19
e.
Number of inferred interactions
238
190
SjS
Sicca
f.
CD46
CLEC
CD45
CD22
NEGR
ESAM
JAM
PARs
ITGB2
ADGRE5
FN1
MK
APP
NOTCH
VCAM
CD99
LAMININ
COLLAGEN
PTN
CXCL
MHC-I
MIF
PDGF
ANGPTL
FGF
MPZ
SELL
IFN-II
BAFF
LCK
PROS
GAS
0.00 0.25 0.50 0.75 1.00
Relative number of interactions
CLEC
CD45
CD22
ADGRE5
ITGB2
CD46
PARs
JAM
ESAM
NEGR
BAFF
LCK
MHC-I
VCAM
CD99
MK
APP
SELL
CXCL
PTN
FN1
NOTCH
MIF
COLLAGEN
LAMININ
IFN-II
MPZ
ANGPTL
PDGF
FGF
PROS
GAS
0.00 0.25 0.50 0.75 1.00
Relative information flow
SjS
Sicca

suppl. fig 4

a.

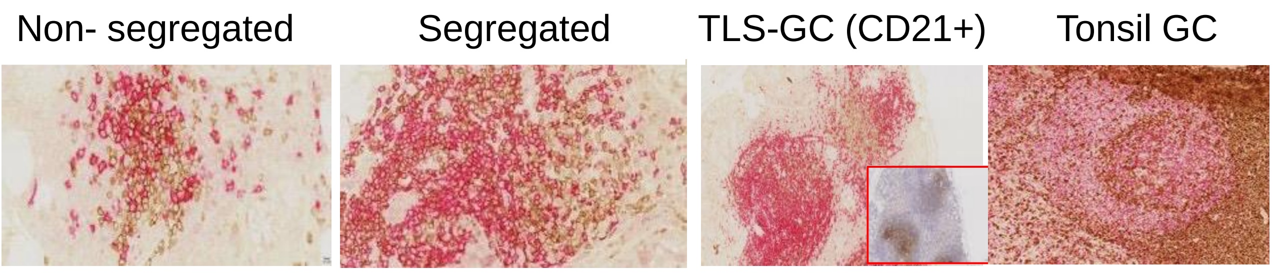


b.

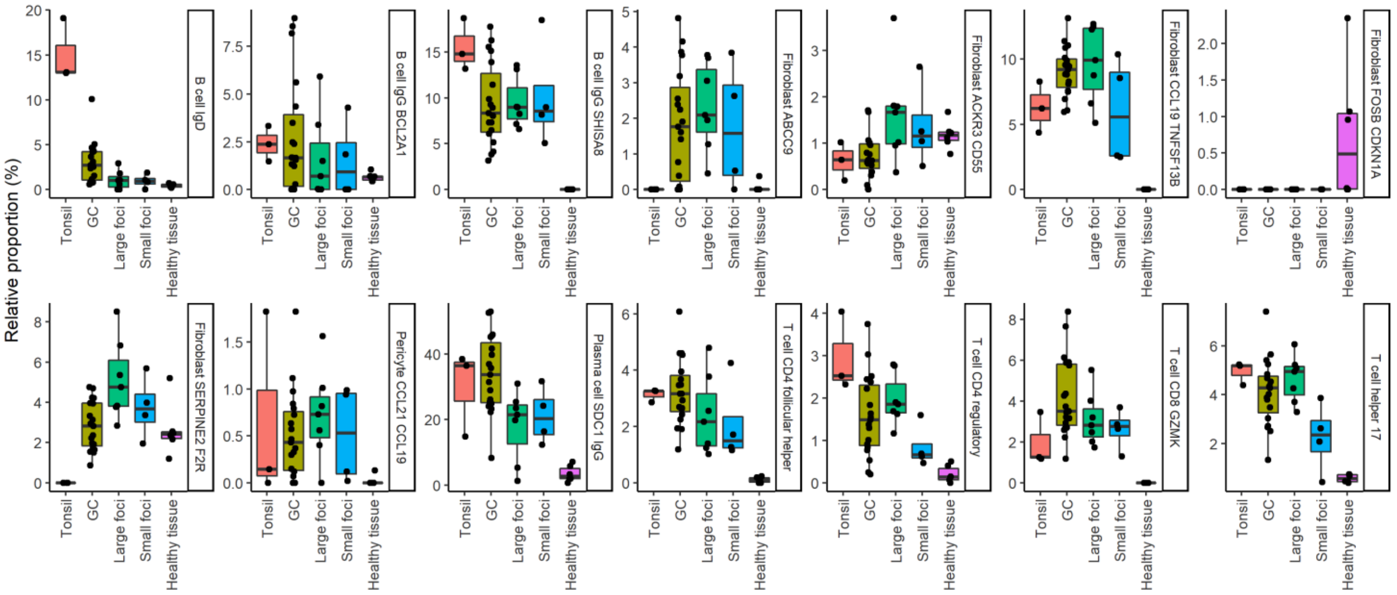


suppl. fig 5

Supplementary figure legends:

**Supplementary figure 1: QC metrics of the SjS 10x data: a,** Expression of marker genes identifying gross-cell identities isolated from Sjogren's minor salivary glands. **b,** Plots of the number of genes, number of UMIs, and percentage mitochondrial genes per sample. **c,** Proportions of gross cell states across samples
